# Chi hotspot control of RecBCD helicase-nuclease by long-range intramolecular signaling

**DOI:** 10.1101/2020.05.04.077495

**Authors:** Susan K. Amundsen, Andrew F. Taylor, Gerald R. Smith

## Abstract

Repair of broken DNA by homologous recombination requires coordinated enzymatic reactions to prepare it for interaction with intact DNA. The multiple activities of enterobacterial RecBCD helicase-nuclease are coordinated by Chi recombination hotspots (5’ GCTGGTGG 3’) recognized during DNA unwinding. Chi is recognized in a tunnel in RecC but activates the RecB nuclease, >25 Ǻ away. How the Chi-dependent signal travels this long distance has been unknown. We found a Chi-recognition-deficient mutant in the RecB helicase domain located >45 Ǻ from both the Chi-recognition site and the nuclease active site. This unexpected observation led us to find additional mutations that reduced or eliminated Chi hotspot activity in each subunit and widely scattered throughout RecBCD. Each mutation alters the intimate contact between one or another pair of subunits in the crystal or cryoEM structures of RecBCD bound to DNA. Collectively, these mutations span a path ∼185 Ǻ long from the Chi recognition site to the nuclease active site. We discuss these surprising results in the context of an intramolecular signal transduction accounting for many previous observations.

## Introduction

Regulation of complex enzymes and reaction pathways is critical for all living organisms, because having too much or too little enzymatic activity, or having spurious activity, can be highly deleterious or lethal. This principle holds true for the repair of broken DNA, which requires in many cases large, complex enzymes. If not properly regulated, DNA repair can fail and produce incomplete or rearranged chromosomes leading to sickness or death. Here, we identify widely scattered positions in the enterobacterial RecBCD helicase-nuclease required for Chi-hotspot regulation of RecBCD and thus proper DNA break repair and genetic recombination.

When DNA is broken, its faithful repair requires interaction with intact homologous DNA, such as the sister chromosome or, in diploids, the homolog. This interaction can produce genetic recombinants if the broken and intact DNAs are genetically different, as often occurs in diploids or when linear DNA is introduced into a cell. In many bacteria, including *Escherichia coli* studied here, the major pathway of homologous recombination and DNA break repair requires RecBCD enzyme, a three subunit, 330 kDa enzyme with DNA unwinding (helicase) and DNA cutting (nuclease) activities (reviewed in ^1^). RecBCD enzymes of enteric bacteria, ranging from *E. coli* to *Proteus mirabilis* and *Vibrio harveyi*, are controlled by a special DNA sequence called Chi (5’ GCTGGTGG 3’) ^2-4^. At Chi the enzyme acts at high frequency, resulting in a high frequency of recombination. Chi is thus a recombination hotspot. After acting at Chi, RecBCD does not act at another Chi site; this regulation assures a single recombination event for each DNA end and regeneration of the circular bacterial chromosome necessary for viability ^5,6^. The molecular mechanism of regulation by Chi has been elucidated progressively, both genetically and biochemically, since its discovery over four decades ago ^1,7,8^. Our results reported here begin to explain at the atomic level Chi’s regulation of RecBCD and may serve as an example by which other complex enzymes are regulated by large conformational changes during their reaction cycle.

The RecBCD reaction cycle begins with RecBCD binding tightly (KD ∼0.1 nM) to the end of broken DNA (Fig. 1A) ^9^. The RecD helicase binds to and moves along the 5’-ended strand rapidly (∼1 kb/sec); the RecB helicase binds to and moves along the 3’-ended strand more slowly (about ½ the rate of RecD) ^10,11^. Consequently, a single-stranded (ss) DNA loop on the 3’-ended strand accumulates, presumably ahead of RecB, and grows with time ^12,13^. The crystal and cryoEM structures of RecBCD bound to DNA show that, when the 3’-ended strand emerges from the RecB helicase, it enters a tunnel in RecC (Fig. 1B) ^14,15^. Mutational analysis indicates that the Chi sequence on the 3’-ended strand is recognized within this RecC tunnel ^16-18^.

**Figure 1.**
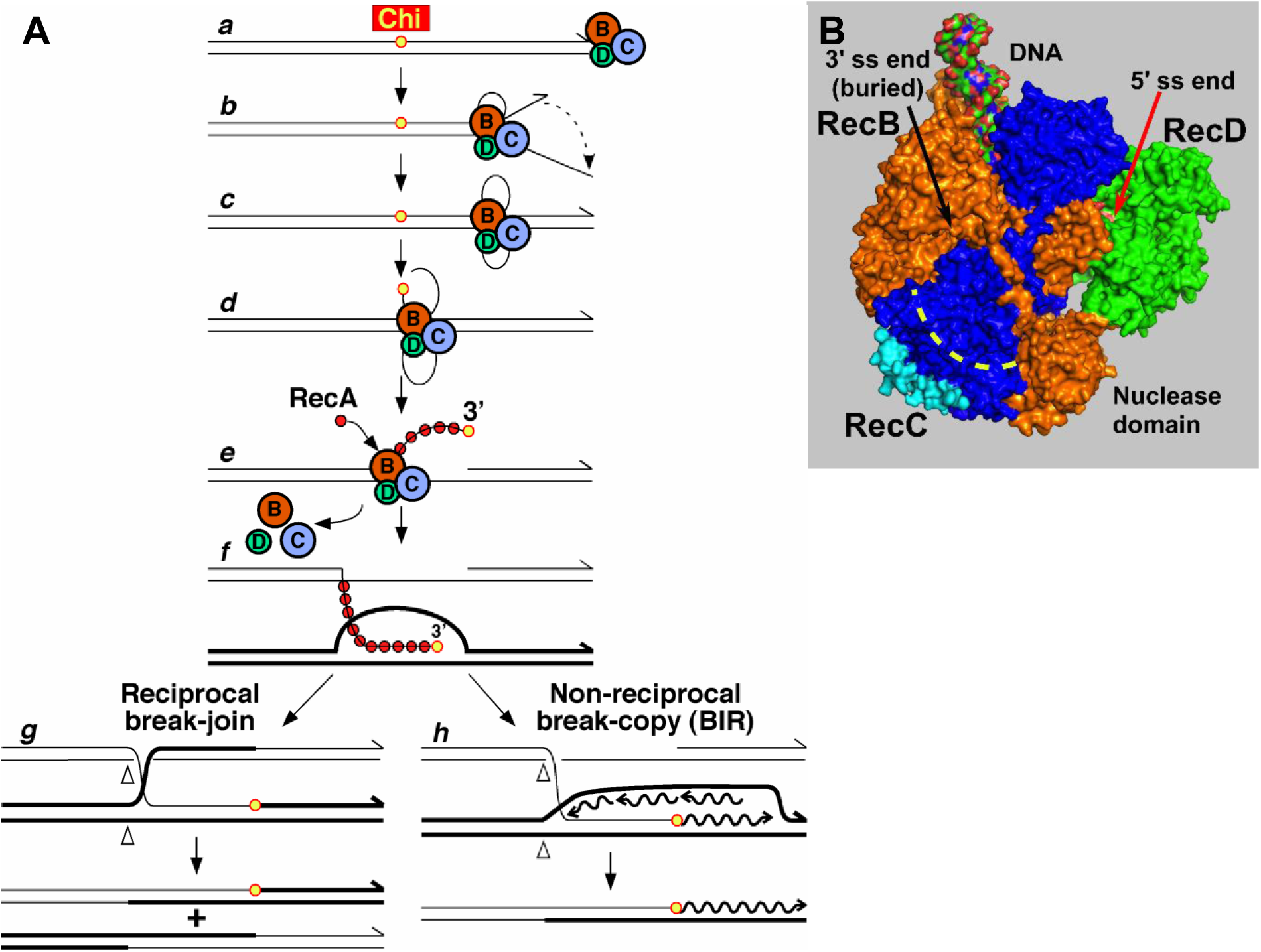
Model for homologous recombination and DNA break repair by RecBCD enzyme and its control by Chi hotspots. (**A**) Pathway of RecBCD-promoted recombination and DNA break repair ^1,23^. RecBCD binds a ds DNA end (***a***) and unwinds the DNA, producing loop-tail structures (***b***) that are converted into twin-loop structures (***c***) by annealing of the tails. At Chi, RecBCD nicks the 3’-ended strand (***d***) and loads RecA (***e***). The ssDNA-RecA filament invades intact homologous DNA to form a D-loop (***f***), which can be converted into a Holliday junction and resolved into reciprocal recombinants (***g***). Alternatively, the 3’-end in the D-loop can prime DNA synthesis and generate a non-reciprocal recombinant (***h***). See ref. ^1^ for discussion of alternative models. (**B**) Atomic structure of RecBCD bound to DNA (PDB 1W36) ^14^. RecB is orange, RecC blue, and RecD green. Yellow dashed line indicates the RecC tunnel in which Chi is recognized. Cyan indicates the RecC patch with differential trypsin-sensitivity during the RecBCD reaction cycle ^19^.

Chi recognition results in multiple changes in the activities of RecBCD ^6,19,20^. Upon encountering Chi, the RecB nuclease cuts the emerging 3’-ended strand a few nucleotides 3’ of the Chi octamer, and the enzyme begins loading RecA DNA strand-exchange protein onto the newly generated 3’-ended strand ^19,21,22^. The RecA-ssDNA filament can invade homologous DNA, and subsequent reactions can produce intact DNA and reciprocal or non-reciprocal recombinants (Fig. 1A) ^1,23^. After cutting at Chi, RecBCD continues to unwind the DNA, but at a slower rate ^24^, as though the slower RecB translocase becomes the leading (unwinding) helicase; the initially faster RecD helicase either stops moving or moves more slowly than before Chi. Although unwinding continues after Chi, the enzyme fails to cut detectably at a subsequently encountered Chi ^6^, as noted above. Furthermore, the three subunits dissociate after cutting at Chi, likely upon leaving the DNA (at the end of linear DNA substrates with purified enzyme) ^20^.

These observations indicate that Chi affects all three subunits of RecBCD. Chi is recognized in a tunnel in RecC but activates the nuclease and RecA-loading activity in RecB; Chi also alters the activity of the RecD helicase. There are additional subunit interactions required for RecBCD’s activities, because *recD* mutants lack Chi hotspot activity and nuclease activity ^25,26^, although the sole nuclease active site is in RecB ^27^. To explore the extent of subunit interactions involved in Chi activity, we searched for mutations outside the RecC tunnel that decrease or abolish Chi hotspot activity. Remarkably, we found mutations with this property at widely scattered positions in all three subunits. We discuss how these positions coordinate RecBCD’s activities and thus regulate DNA break repair and recombination.

## Results

### A recombination-proficient *recB* mutant with reduced Chi hotspot activity

A search for *E. coli* mutants that excise transposon *Tn10* at a high rate yielded mutations in many genes, including, unexpectedly, two that complemented as a *recC* or *recB* mutation ^28^. Later research showed that one, *recC343*, changes proline 666 to leucine (P666L; 5’ CCA 3’ changed to 5’ CTA 3’), which is on the surface of the RecC tunnel in which Chi is recognized ^14,29^. Dozens of other RecC tunnel mutants have the same or similar phenotype – little or no Chi hotspot activity but retention of recombination-proficiency and “general” (Chi-independent) nuclease activity ^16-18,30^. More surprising, however, was *recB344*: this mutant has significantly reduced Chi hotspot activity, the ratio of recombinant frequency in a genetic interval with Chi to that in the same interval without Chi ^31^. Chi activity was 2.1 ± 0.033 in *recB344* but 4.9 ± 0.021 in concurrent wt crosses (1 indicates no Chi activity; data are mean ± SEM with n = 3) (Table 1) (see also ^18,28^). It also had reduced cutting of DNA at Chi (Fig. S1), as expected from the genetic assays. The *recB344* mutant retains, however, recombination-proficiency in *E. coli* Hfr and phage lambda crosses (93% and 94%, respectively, of wt) and general nuclease activity (60% of wt) (Tables 1 and 2) ^18,28^. Thus, it has the same overall phenotype as RecC tunnel mutants with reduced Chi hotspot activity.

**Table 1.**
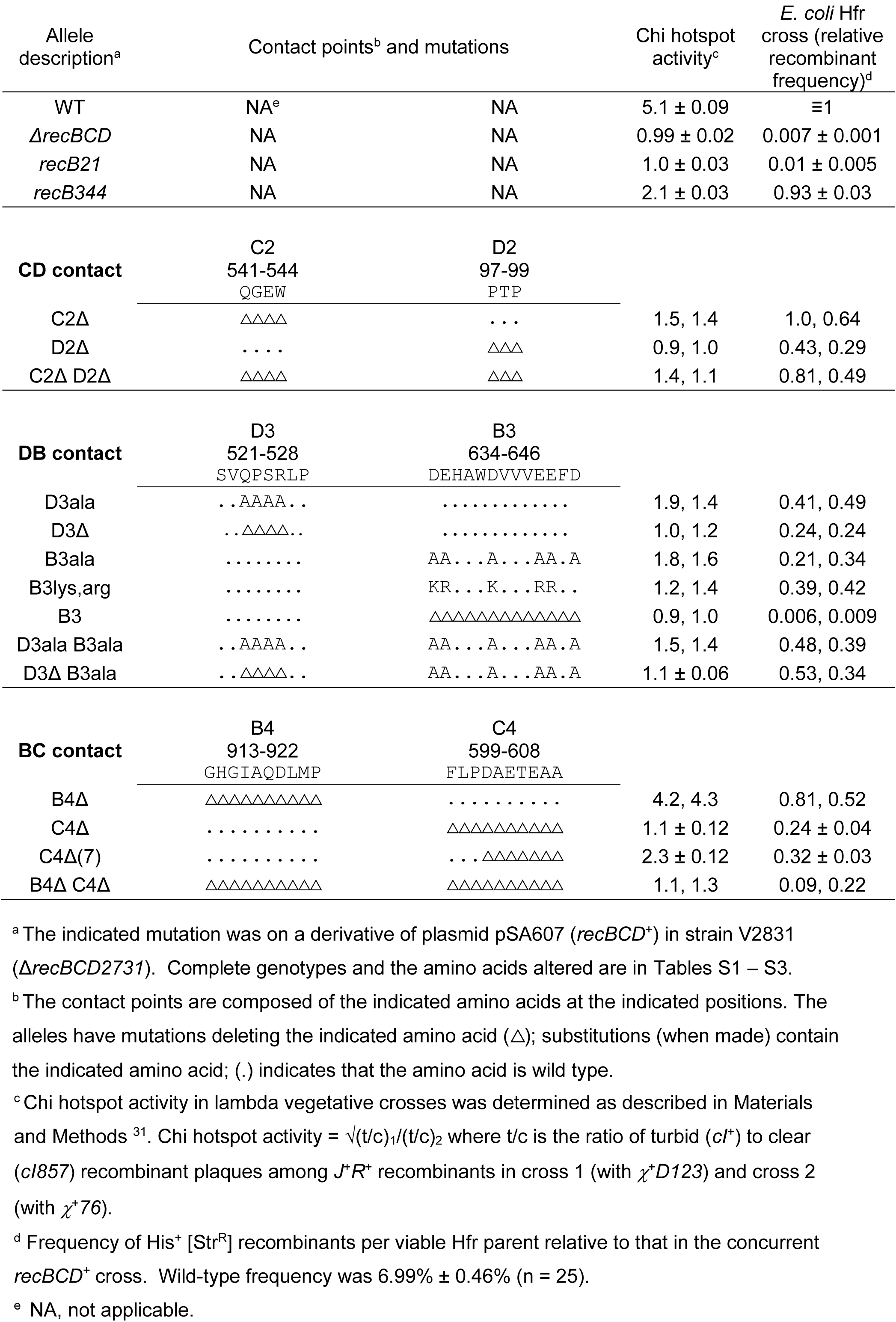
Mutants altered in RecC-RecD contact (CD), RecD-RecB contact (DB), and RecB-RecC contact (BC) have little or no Chi hotspot activity

**Table 2.** RecBCD contact mutant enzymatic and genetic activities.

| Con-<br>tact<br>point | Allele<br>description<sup>a</sup> | ATP-dep. nuclease<br>activity<br>(units/mg protein)<sup>b</sup> |  | T4 2<sup>-</sup><br>eop<sup>c</sup> | DNA<br>unwinding<sup>d</sup> | Chi<br>cutting<sup>d</sup> | Chi hotspot<br>activity<sup>e</sup> |
| --- | --- | --- | --- | --- | --- | --- | --- |
| | | 25 μM<br>ATP | 1 mM<br>ATP | | | | |
| | *recBCD+* | 440, 250 | 280, 280 | $<2 \times 10^{-8}$ | + | + | $5.1 \pm 0.09$ |
| | Δ*recBCD* | $<2$ | ND<sup>f</sup> | 1.0 | - | - | $0.99 \pm 0.02$ |
| | *recB344* | 220 | ND | ND | + | ± | $2.1 \pm 0.03$ |
| **CD** | C2Δ | 780 | 360 | $<2 \times 10^{-8}$ | + | - | 1.5, 1.4 |
| | D2Δ | 3.5 | $<2$ | $1.6 \times 10^{-3}$ | + | - | 0.9, 1.0 |
| | C2Δ D2Δ | 10 | 16 | $1.3 \times 10^{-3}$ | + | - | 1.4, 1.1 |
| | C2ala | 5.2 | 12 | $<2 \times 10^{-8}$ | + | - | 3.2, 3.5 |
| | D2ala | 8.7 | 21 | $<2 \times 10^{-8}$ | + | - | 2.4, 2.9 |
| | C2ala D2ala | 1.7 | 1.7 | $<2 \times 10^{-8}$ | + | - | 2.2, 2.3 |
| **DB** | B3Δ | 3.5 | 23 | $5.1 \times 10^{-4}$ | + | - | 0.9, 1.0 |
| | B3ala | 400, 610 | 300, 360 | $<2 \times 10^{-8}$ | + | + | 1.8, 1.6 |
| | D3ala | 310, 470 | 340, 330 | $<2 \times 10^{-8}$ | + | + | 1.9, 1.4 |
| | B3ala D3ala | 37, 37 | 96, 93 | $<2 \times 10^{-8}$ | + | - | 1.5, 1.4 |
| | D3Δ | 14, 38 | 68, 68 | $3.1 \times 10^{-6}$ | + | - | 1.0, 1.2 |
| | B3Δ(10) | 400, 370 | 240, 240 | $<2 \times 10^{-8}$ | + | + | 4.6, 4.2 |
| | B3ala D3Δ | 12, 16 | 47, 35 | $4.6 \times 10^{-4}$ | + | - | $1.1 \pm 0.06$ |
| **BC** | B4Δ | 700 | 350 | $<2 \times 10^{-8}$ | + | + | 4.2, 4.3 |
| | C4Δ | 10 | 23 | $<2 \times 10^{-8}$ | + | - | $1.1 \pm 0.12$ |
| | B4Δ C4Δ | 12 | 16 | $1.9 \times 10^{-3}$ | + | - | 1.1, 1.3 |

<sup>a</sup> The indicated mutation was on a derivative of plasmid pSA607 (*recBC*<sup>2773</sup>*D*) in strain V2831 ( $\Delta$ *recBCD2731*). Alleles are described in Tables 1 and S1 – S3.
<sup>b</sup> Specific activity in units of ATP-dependent ds DNA nuclease/mg of protein at 25 μM ATP or 1 mM ATP. ND, not determined.
<sup>c</sup> Efficiency of plating (eop) is the phage titer on the indicated strain divided by the titer on strain V2831 ( $\Delta$ *recBCD2731*).
<sup>d</sup> Activity in extracts (Figs. 4 and S1). +, activity present. -, activity absent.
<sup>e</sup> Data from Tables 1 and S1 – S3.
<sup>f</sup> ND, not determined.

Also surprising was the location of the *recB344* mutation. DNA sequencing showed that it changes 5’ GCC 3’ to 5’ GTC 3’, which changes alanine 64 to valine (A64V). A64 is part of an α-helix within the RecB helicase domain ^14,29^. In the cryoEM structure in which the Chi octamer is in the RecC tunnel, A64 is 24 – 42 Ǻ from the Chi nucleotides and 65 Ǻ from the RecB nuclease active site ^15^. These results show that positions in RecBCD far from either the site of Chi recognition (in the RecC tunnel) or the nuclease active site (in the RecB nuclease domain) are involved in Chi hotspot activity. They also led us to seek additional RecBCD mutants with little or no Chi hotspot activity but with amino acid alterations far from the RecC tunnel or the RecB nuclease active site.

### Search for additional wide-spread mutations with the Chi-recognition-deficient phenotype

The recognition of Chi in the RecC tunnel but its stimulation of the RecB nuclease indicates that some “signal” must be transmitted through the enzyme from the tunnel to the nuclease. A search for a Chi-dependent covalent modification, such as phosphorylation, that could account for the Chi-dependent alteration of RecBCD activities was negative. Instead, we found evidence for a Chi-dependent conformational change: a patch on the RecC surface (Fig. 1B) is sensitive to four proteases before DNA is bound and becomes markedly more resistant when DNA is bound ^19^. During unwinding this patch remains resistant up to Chi but becomes protease-sensitive again after Chi’s encounter. The *recB344* mutation may affect this Chi-dependent conformational change. Other mutations could have a similar effect, by blocking a conformational change from one subunit to another. If so, we reasoned that points of intimate contact between subunits might well be involved in the response to Chi.

To test this hypothesis, we examined crystal and cryoEM structures of RecBCD ^14,15,29^ for points of contact between RecC and RecD (called “CD”), between RecD and RecB (“DB”), and in or near the RecB nuclease domain close to the exit of the RecC tunnel (“BC”) (Fig. 2). (To aid Discussion, we designate each side of the contact as C2 D2, D3 B3, and B4 C4, respectively; Fig. 2.) Contact points for DB and BC were discussed by Wilkinson et al. ^29^ (see Discussion). We then mutated, by deletion or amino acid substitutions, these points of contact and assayed the genetic and enzymatic activities as for *recB344* and other Chi-recognition mutants. (Note that there is no direct assay for Chi recognition, such as binding of Chi DNA to RecBCD, since Chi is recognized only during active unwinding by RecBCD; consequently, only the effect of Chi can be assayed, such as cutting of DNA at Chi and Chi genetic hotspot activity.) As described below, we isolated multiple mutants altered at each contact point. Many of these mutants have reduced or undetectable Chi hotspot and cutting activities, like previously described Chi recognition mutants ^16-18,30^.

**Figure 2.**
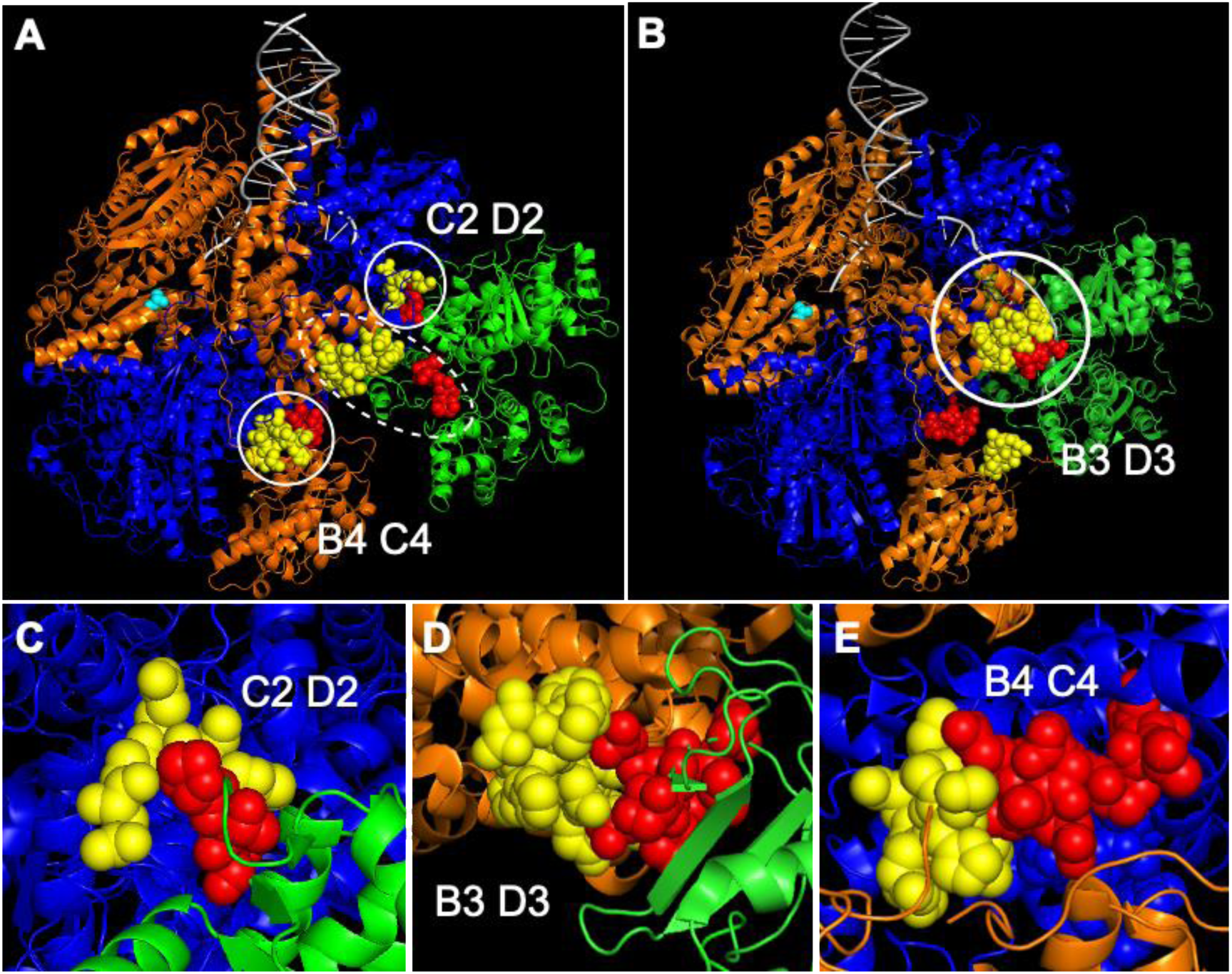
Contact points between RecBCD subunits tested for a role in Chi hotspot activity. Atomic structures of RecBCD showing contacts between RecB (orange), RecC (blue), and RecD (green); DNA is grey. Points of contact studied here are represented as spheres; the rest of the molecule is represented as cartoons (PyMol version 2.2.0). Note that the RecB-RecD contact points are separated by >20 Ǻ in the crystal structure (dotted circle in **A**) PDB 1W36 ^14^ but are close to each other in the cryoEM structure (solid circle in **B**) PDB 5LD2 ^15,29^. Note that, conversely, a domain of ∼70 amino acids in RecD is ordered in the cryoEM structures but disordered in the crystal structures. (**C**) Contact point CD. C2 (amino acids QGEW at positions 541 – 544 of RecC) is yellow, and D2 (PTP at positions 97 – 99 of RecD) is red. Shown is part of the crystal structure PDB 1W36. (**D**) Contact point DB. B3 (DEHAWDVVVEEFD at positions 634 – 646 of RecB) is yellow, and D3 (amino acids SVQPSRLP at positions 521 – 528 of RecD) is red. Shown is part of the cryoEM structure PDB 5LD2. (**E**) Contact point BC. B4 (amino acids GHGIAQDLMP at positions 913 – 922 of RecB) is yellow, and C4 (amino acids FLPDAETEAA at positions 599 – 608 of RecC) is red. Shown is part of the crystal structure PDB 1W36.

### RecC-RecD interaction

In the published crystal and cryoEM structures (Figs. 2A and 2B) ^14,15,29^, four contiguous amino acids (here named C2) in RecC make intimate contact with three contiguous amino acids (named D2) in RecD (Fig. 2C; Table 1). Deletion of C2 (C2Δ) strongly reduced Chi hotspot activity to 1.5 (from 5.1 in wild type), and deletion of D2 (D2Δ) abolished Chi hotspot activity (Chi activity = 1.0) (Table 1). As expected, the double mutant C2Δ D2Δ also had no significant Chi activity. Substitution of alanine for the amino acids of C2 or D2 (C2ala or D2ala) also significantly reduced Chi activity, to 3.3 or 2.7, respectively (Table S1). These mutants had slightly reduced recombination proficiency in Hfr matings (∼40 – 80% of wt); a *recBCD* null mutant’s proficiency is much less (0.7% of wt; Tables 1 and S1; Fig. 3B). Thus, Chi hotspot activity requires the point of contact (CD) between RecC and RecD.

**Figure 3.**
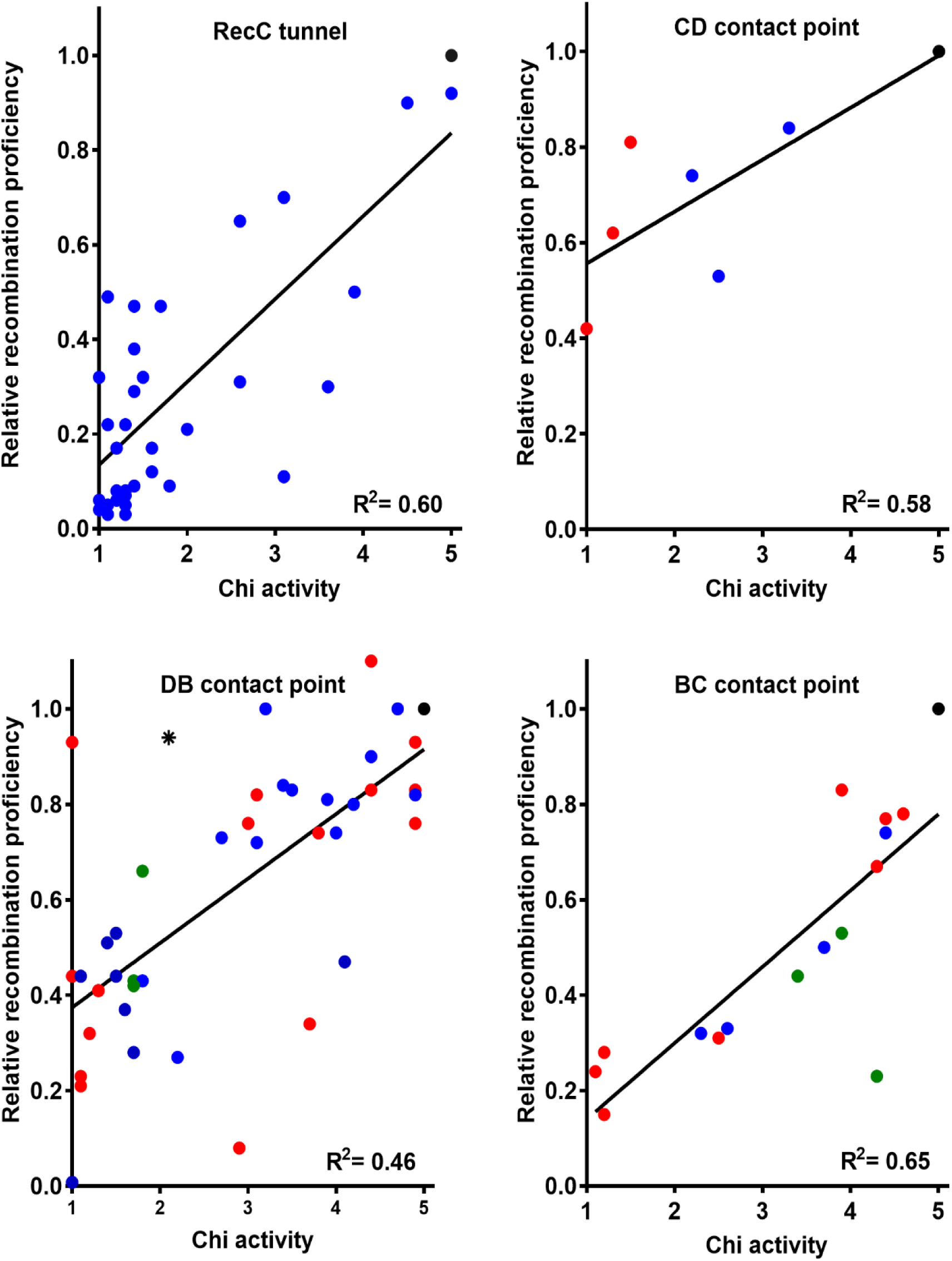
*E. coli* Hfr recombination proficiency is positively correlated with Chi hotspot activity. Red data points are for deletion mutants, blue for substitution mutants, and green for mutants with a substitution and deletion; black point is wild type, and star is *recB344*. Linear regression lines and the coefficient of determination (R^2^) are shown. Data are from Tables 1, 2, S1, S2, and S3 and refs ^16-18^.

### RecD-RecB interaction

In the cryoEM structures of RecBCD bound to ds DNA ^15,29^, four contiguous amino acids with overall positive charge (QPSR in the center of D3) in RecD contact an α-helix of 13 amino acids (B3), six with negative charge (D or E), in RecB (Fig. 2D; Table 1). As for the CD interaction, complete deletion of B3 or the central part of D3 eliminated Chi hotspot activity (Table 1). Substitution of alanine for the central four amino acids in D3 or for the six negative amino acids in B3 or both sets strongly reduced Chi activity (to 1.7, 1.7, and 1.5, respectively; Table 1). A second double mutation (B3ala D3Δ) eliminated Chi activity (1.1). Hfr recombination proficiency was reduced to 20 – 50% of the wild-type value, except in the B3Δ single mutant, which had recombination proficiency similar to that of a *recBCD* null mutant (Table 1). These results show that the DB point of contact is essential for Chi activity.

As noted above, deletion of the central four amino acids in D3 (QPSR) eliminated Chi activity. To determine which of these four amino acids is critical, we deleted separately the left two (QP) and the right two (SR) amino acids. Surprisingly, both mutants had nearly full Chi activity (4.4 and 5.0, respectively; Table S2A). We then deleted other combinations of amino acids in D3 (SVQPSRLP). Some of these deletions eliminated Chi activity (e.g., deletion of SV, deletion of QP plus LP, or deletion of SRLP); other deletions strongly reduced Chi activity (e.g., deletion of just V, to 1.5). These results show that D3 and its surround are critical for Chi activity and suggest a more complex effect than simple deletion of a few amino acids as the basis for loss of Chi activity. Indeed, except in one case, substitution of alanine for one to four of the amino acids in SVQPSRLP also significantly reduced Chi activity (to 1.4 – 3.5).

We similarly explored the requirements for amino acids in B3 (DEHAWDVVVEEFD). Deletion of only 3, 5, 8, or 10 amino acids, collectively spanning all points of the 13 amino acids deleted in ΔB3, left significant but reduced Chi activity (2.9 – 4.4; Table S2B), suggesting that no individual part of B3 is essential. Four or six alanine substitutions distributed throughout B3 substantially reduced Chi activity (to ∼2.0). Other substitutions in B3 also significantly reduced Chi activity (e.g., ………E..D to ………Q..S had Chi activity of 3.4, and .R…N…QK.S had Chi activity of 3.0. Substitutions of only two D or E residues with K or R left nearly full Chi activity (4.8 and 4.1), but similar K and R substitutions of four or five residues nearly eliminated Chi activity (1.8, 1.6, and 1.3). Thus, both the presence and the amino-acid composition of B3 are critical for Chi activity.

Double B3 D3 mutants were similar to the stronger single mutant, except in one case. An additive effect of .R…N…QK.S in B3 and ..E.DE.. in D3 was observed in the double mutant: Chi activities were 3.0, 3.2, and 1.8, respectively (Table S2C). Thus, the role of DB is more complex than that of CD, but nevertheless this couple also plays an essential role in Chi hotspot activity.

### RecB-RecC interaction

In the published crystal structures of RecBCD ^14,32^, a 10-amino-acid α-helix (B4) in the RecB nuclease domain is close to a 25-amino-acid α-helix in RecC; we define C4 as the seven amino acids at the N-terminus of this RecC α-helix plus the adjacent three amino acids, which form a turn (Fig. 2E; Table 1). The B4 α-helix has been postulated to control RecBCD nuclease activity ^14,29^, but deletion of B4 only marginally reduced Chi activity (to 4.3) and had no significant effect on Chi-independent nuclease activity (Tables 1 and 2). Deletion of C4, however, eliminated Chi activity, as did the double deletion (B4Δ C4Δ) (Table 1). Deletion of seven amino acids in C4 also strongly reduced Chi activity (to 2.3) (Table 1). Alanine substitutions of the ten amino acids in B4 or three of the ten amino acids in C4 slightly reduced Chi activity (to 3.7 and 4.4, respectively; Table S3). These results show that at least part of C4 is essential for Chi activity but that B4 plays at most a minor role on its own.

We noted that the B4 and C4 α-helices ordered in the crystal structure are close to a seven-amino-acid loop of RecD (D4, amino acids 469 – 475; Table 1) in the cryoEM structures (PDB 5LD2, 6SJB, 6SJE, 6SJF, 6SJG, 6T2U, and 6T2V; Figs. 2B and S2) ^15,29^. This loop in RecD and 50 surrounding amino acids are disordered in both molecules in the asymmetric unit of both crystal structures (PDB 1W36 and 3K70; Fig. 2A) ^14,32^, and B4 is disordered in the cryoEM structures. This outcome indicates that B4 and the loop in RecD can adopt alternative structures. Deletion of this loop or changing each amino acid to alanine had little if any effect on Chi activity (3.9 or 4.4) or recombination potential (74 or 83% of wild type) (Table S3). Control of RecBCD thus appears to be largely independent of this loop in RecD.

### Recombination proficiency is positively correlated with Chi hotspot activity

The critical role of Chi in regulating RecBCD’s activities predicts that lowering the Chi hotspot activity would also lower recombination proficiency. We tested this hypothesis by plotting the two values for each of the mutants discussed above [RecC tunnel and contact points CD, DB (with *recB344*), and BC] (Fig. 3). The correlation was significantly positive (R^2^ = 0.60, 0.58, 0.46, and 0.65 for these sets, respectively; p = <0.0001, 0.047, <0.0001, and <0.0002, respectively). We noted that the intercept of Hfr recombination proficiency at no Chi hotspot activity (0.27; mean of the four values in Fig. 3) was comparable to the ratio (0.20) of lambda recombination without and with Chi in the Chi hotspot crosses. In other words, removing a Chi site reduces lambda recombination to about the same extent that inactivating any of the four contact points reduces Hfr recombination. Residual recombination reflects RecBCD’s innate ability to promote recombination without Chi or the signal it elicits.

### New mutants have DNA unwinding and general nuclease activity but have lost Chi cutting

To test more directly the effect of the mutations on RecBCD activities, we assayed DNA unwinding, cutting of DNA at Chi, and general nuclease activities. Representative mutants altered at each of the contact points were tested. In some cases, the mutants retained approximately wild-type levels of DNA unwinding activity (Figs. 4 and S1), but in other mutants there was significant reduction. As expected, Chi cutting was observed in mutants with full or slightly reduced Chi hotspot activity (>4.3) but not in mutants with little or no Chi activity (<1.6). Chi cutting was not observed in CD mutants with partial Chi genetic activity (2.2 – 3.4) but was observed in two DB mutants (B3ala and D3ala) with low Chi hotspot activity (1.7 – 1.8) (Table 2). The difference may reflect intracellular *vs.* extracellular conditions for the assays.

**Figure 4.**
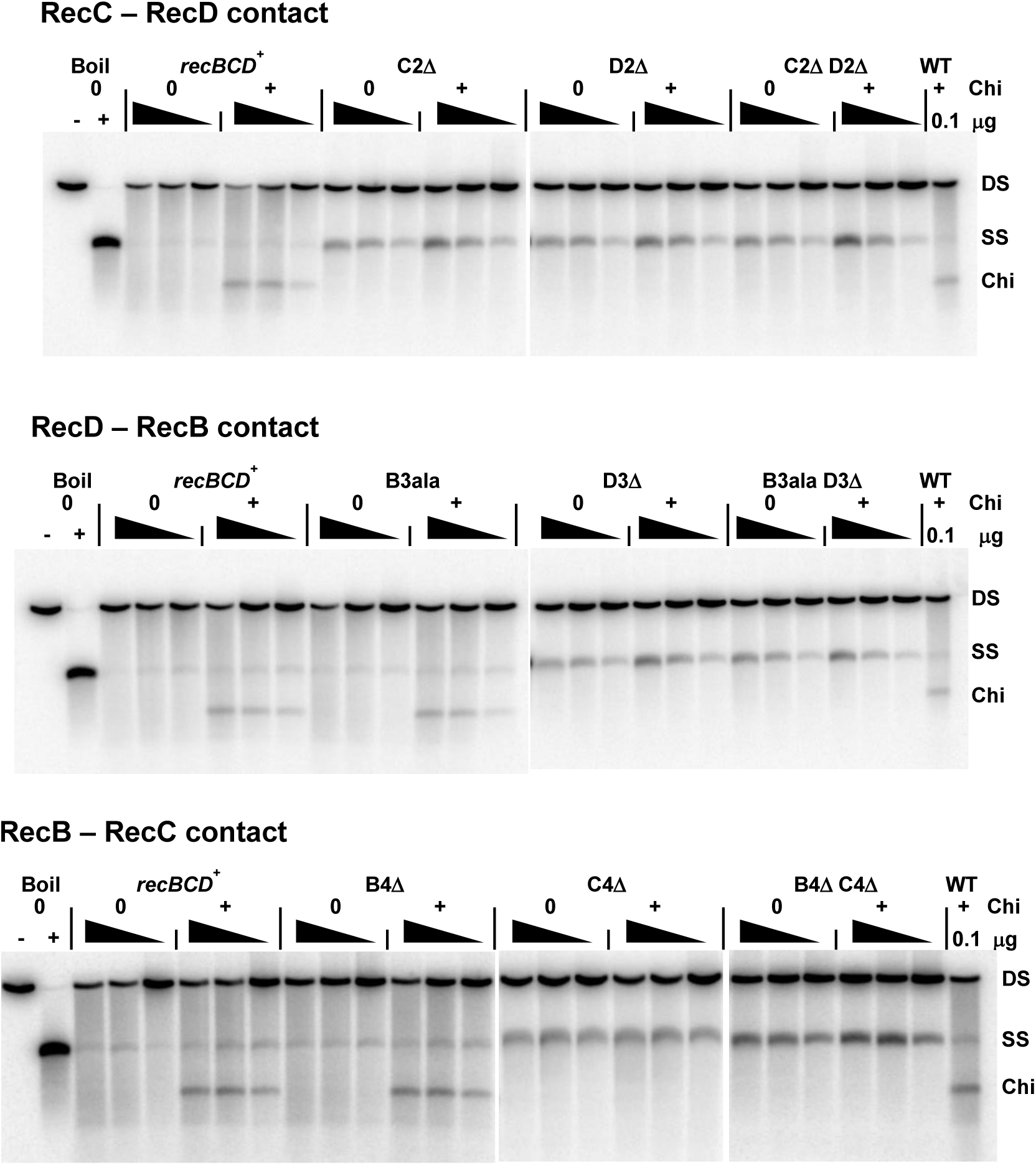
RecBCD contact-point mutants retain DNA unwinding activity but have reduced or undetectable cutting of DNA at Chi hotspots. Extracts of *recBCD*^+^ (wild type or WT; 0.3, 0.1 or 0.03 μg protein) and the indicated mutants (1, 0.3 or 0.1 μg protein) were assayed for unwinding and cutting of linear pBR322 DNA (4.3 kb long) with or without a Chi site (*χ*^+^*F225*) 1470 bp from the 5’ [^32^P]-labelled DNA end. Note that the Chi-cut species and ss DNA are reaction intermediates and their observed amount is not necessarily a linear function of enzyme amount. ds substrate (DS), unwound ss DNA (SS), and Chi-cut DNA (Chi) are indicated.

Chi-independent nuclease activity was nearly wild-type in some mutants but was low in others (Table 2). In most of the latter cases, nuclease activity was about 2-fold higher with 1 mM ATP than with 25 µM ATP; wild-type RecBCD activity, by contrast, was about 2-fold <u>lower</u> with 1 mM ATP, as reported previously ^33^. Possible explanations for this phenotype are in the Discussion. All of the mutants studied here do, however, block the growth of T4 *gene 2* mutants, showing that they retain nuclease activity in living cells ^34^.

## Discussion

The results reported here greatly expand the set of mutants with reduced control of RecBCD enzyme by Chi hotspots of recombination. They reveal that points of contact between each of the enzyme’s three subunits are essential for Chi activity (Fig. 2C – E), and they provide important material for more direct biophysical assays of the molecular mechanism of this control.

The 63 mutants studied here have a wide range of effects on Chi activity, from no significant effect to complete loss of Chi activity. At each of the three tested points of intersubunit contact, we found mutants that lacked Chi activity. For two of these contact points (CD and DB), we found such mutations in each of the two interacting subunits (RecC-RecD and RecD-RecB, respectively). For the other contact point (BC), mutations in RecC abolished Chi activity, but none of the RecB or RecD mutants tested dramatically lowered Chi activity. Because the nuclease active site is in RecB ^35^ and a part of RecC essential for Chi activity (C4 in Table 1 and Fig. 2E) is close to the RecB nuclease domain, it seems likely that RecC and RecB must contact each other to effect Chi hotspot activity. Thus, our choosing points of contact to mutate, based on the crystal and cryoEM structures, successfully identified areas of RecBCD far from the Chi-recognition and nuclease sites critical for Chi’s control of RecBCD.

It is noteworthy that two of the points of contact are positioned differently in the crystal and cryoEM structures (Figs. 2A and 2B) ^14,15^. The RecD-RecB contact points intimately interact in the cryoEM structures (Fig. 2B and 2D) but are >15 Ǻ apart in the crystal structures (Fig. 2A). Conversely, the RecC-RecB contact points intimately interact in the crystal structures (Fig. 2A and 2E) but are >20 Ǻ apart in the cryoEM structures (Fig. 2B). This is evidence that these points can move relative to each other, in accord with conformational changes of RecBCD being important for Chi hotspot activity. In addition, one of the proposed contact points (B4) is ordered in the crystal structure but not in the cryoEM structures. This outcome also indicates flexibility in contact points, as expected for parts of the protein that interact differently during steps of the reaction cycle.

Based on the crystal and cryoEM structures of RecBCD bound to a set of forked DNA substrates with increasingly long 5’ single-stranded DNA extensions, Wigley’s group has discussed the roles of contact points DB and BC (Fig. 2D and 2E). When the 5’ extension of the DNA is elongated from four to ten nucleotides, parts of RecD become more ordered ^14,32^. An additional two nucleotides on the 5’ extension result in larger changes ^29^. The SH3-like domain of RecD becomes well-ordered, and contact point D3, in the SH3 domain, touches contact point B3. There is also a major change in the B4 α-helix. The C4 α-helix moves a few Ǻ and apparently allows the B4 α-helix to become disordered and to move away from the RecB nuclease active site. They postulated that this movement of ∼15 Ǻ of what they termed the RecB “linker,” which contains B4, was necessary to convert RecBCD from an inactive nuclease (when bound to a ds DNA end) into an active nuclease (during unwinding). Deletion of B4, however, had little effect on RecBCD nuclease or Chi hotspot activity (Tables 1 and 2; Fig. 2E).

Comparison of cryoEM structures of RecBCD bound to a 3’ extension without Chi or with Chi appropriately placed for its recognition by the RecC tunnel revealed additional small changes in the position and orientation of the RecB nuclease domain ^15^. These changes were postulated to inactivate the nuclease after making its final cleavage, on the 3’-ended strand upon encountering Chi, a property of purified RecBCD under reaction conditions with high [Mg^2+^] but not, the evidence indicates, in living cells ^1,36,37^. With high [Mg^2+^], purified RecBCD begins to cut the 5’-ended strand at Chi ^38,39^, but the basis for this switch in strand cutting was not addressed by Cheng et al. ^15^. The predictable consequences of these structural changes were not tested, for example by the type of mutational analysis used here. Our data here demonstrate the importance of the points of contact in the structures reported by Wigley’s group.

A quarter of the mutations altering the contact points (16 of 63 analyzed here) retained nearly full Chi hotspot activity (4.1 – 5.0), recombination proficiency, and the enzymatic activities tested (Tables 1, 2, and S1 – S3; Figs. 4 and S1). Thus, some alterations as great as deletion of ten amino acids [B3Δ(10) in RecB at the DB contact point (Tables 2 and S2) had little effect. This outcome shows that not all structural alterations, even large ones, block Chi’s control of RecBCD and its other activities. It also supports the view that the contact points in mutants that do have an effect have a direct role in Chi activity as proposed in the models discussed below (Fig. 5). Additional points in RecBCD, such as RecB A64 in the helicase domain and changed to V64 in the *recB344* mutant (Table 1 and 2), are also likely critical for Chi activity, but more complex analyses, such as molecular dynamics, may be required to reliably predict additional points to mutate.

**Fig. 5.**
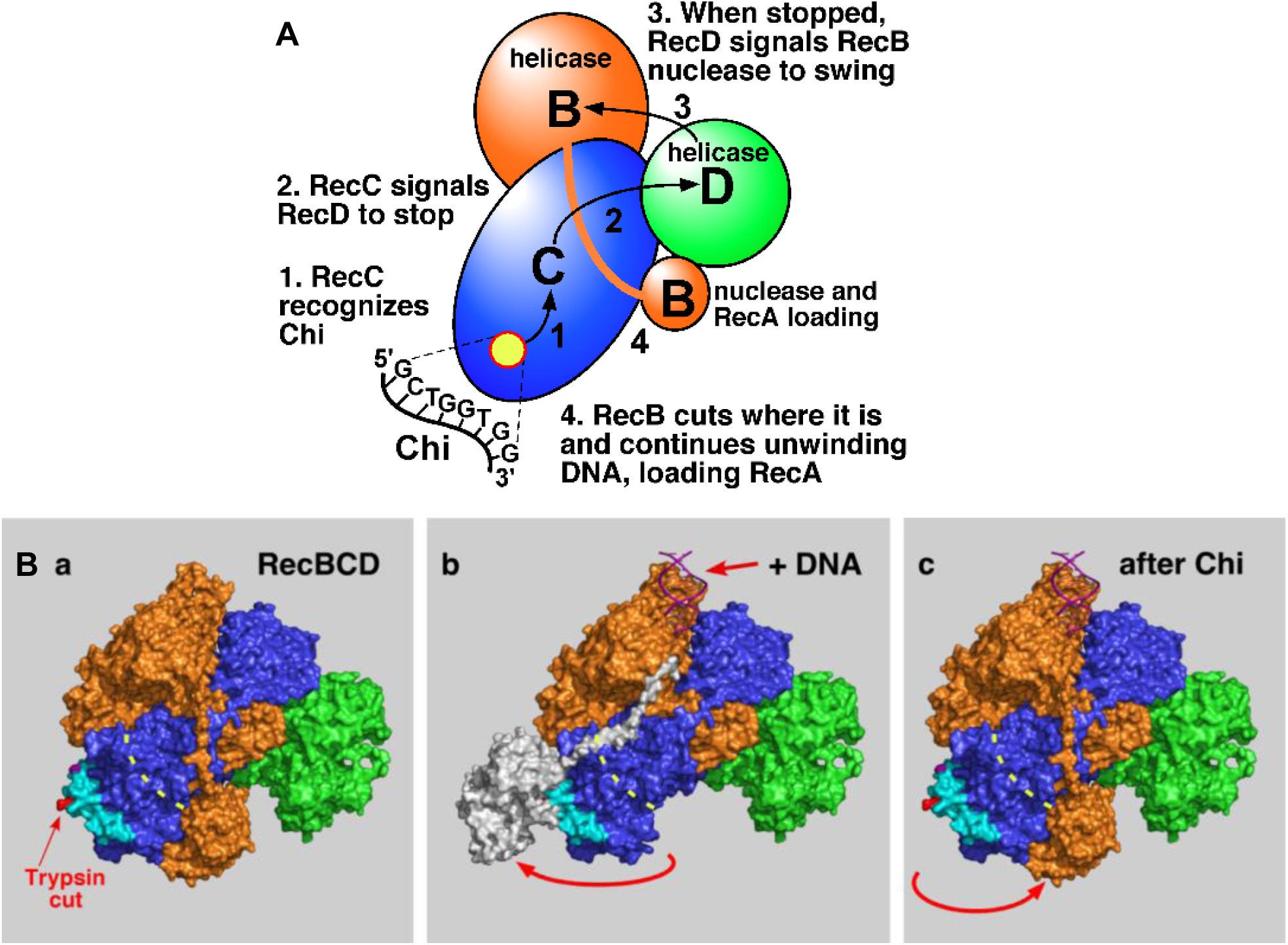
Models for Chi hotspot control of RecBCD enzyme. (**A**) Signal transduction model for RecBCD regulation by Chi ^41^. (**B**) Nuclease swing model for Chi’s control of RecBCD ^19^. Cyan indicates the RecC patch that is differentially sensitive to proteases ^19^. Grey indicates the RecB nuclease domain and the tether connecting it to the RecB helicase domain; it was positioned by hand in the middle panel (**b**). (**a**) Before DNA is bound. (**b**) After DNA is bound but before Chi is encountered. (**c**) After Chi is encountered during unwinding. Modified from ^19^.

Here and elsewhere, we have identified five points in RecBCD enzyme at which mutations reduce or abolish Chi activity. These include mutations in the RecC tunnel (Chi recognition) ^16-18^ and in each of the three contact points studied here (CD, DB, and BC) (Tables 1 and S1 – S3; Figs. 1 and 2). Mutations in the RecB tether connecting the RecB helicase and nuclease domains show that the tether has to be long enough and sufficiently flexible to allow the nuclease domain to swing and act at Chi (Figs. 5B and S3) ^40^. Each of these pairs of contact points spans a distance as great as 80 Ǻ; the maximal distance between points in RecBCD is ∼140 Ǻ. The sum of distances from Chi recognition in the RecC tunnel to point CD, then to point DB, then to point BC, and finally to the nuclease active site is ∼185 Ǻ. The signal from Chi to the nuclease domain thus traverses much of the expanse of the enzyme even though in the reported structures the Chi recognition site is only ∼25 Ǻ from the nuclease active site. This result indicates that a high degree of coordination among the three subunits is required to control the numerous activities of RecBCD.

We have previously proposed a signal transduction model for Chi’s control of RecBCD’s multiple activities (Fig. 5A) ^41^ and a more detailed model (nuclease-swing) for one step (Fig. 5B) ^19^. In these models, Chi is recognized in the RecC tunnel (step 1); RecC then signals RecD helicase to stop (step 2). When RecD is stopped, it signals the RecB nuclease domain to swing from its inactive position on the “left” side of RecC to the RecC tunnel exit (step 3). The nuclease then cuts the DNA a few nucleotides 3’ of the Chi octamer and begins loading RecA strand-exchange protein onto the newly generated 3’ end (step 4). The mutants described here are consistent with step 2 (CD mutants), step 3 (DB mutants), and step 4 (BC mutants) being disrupted. Alteration of each contact point reduces or eliminates the end result of this “pathway” – Chi hotspot-stimulated recombination. The properties of dozens of mutants in the RecC tunnel and the RecB tether also support these models.

The mutants described here provide excellent material for biophysical tests of these models. Possible methods include Förster resonance energy transfer (FRET). RecBCD enzyme with fluorescent moieties at two points, for example on the nuclease domain and at the RecC tunnel exit, could show that the nuclease domain swings as inferred from previous studies (Fig. 5B) ^19^. Such structural changes might also be visible by cryoEM of RecBCD stopped during active unwinding before and after Chi. Single-molecule unwinding experiments, such as those using DNA curtains ^42^, could also show pausing at Chi or not. These methods require advanced equipment not currently available for our studies.

Some mutants studied here unexpectedly had lower “general” (ATP-dependent but Chi-independent) nuclease activity than the wild type (Table 2). These mutants’ nuclease activities were higher with 1 mM ATP than with 0.025 mM ATP, the concentration in the standard nuclease assay ^33^, whereas the wild-type nuclease activity is <u>lower</u> with 1 mM ATP, which is more nearly physiological. In the nuclease-swing model (Fig. 5B), the nuclease domain can be positioned at the exit of the RecC tunnel, where it can cut DNA, or on the protease-protected patch of RecC (Figs. 1B and 5B), where it cannot cut DNA ^19^. We propose that in these mutants the nuclease domain remains, during unwinding, more often than in wild type at the protease-sensitive patch on RecC and thus far from the RecC tunnel exit and unable to cut DNA frequently (Fig. 5B). In this view, the activity of the nuclease reflects the fraction of time the nuclease domain resides at the tunnel exit. This fraction may be altered by the mutations studied here and by the ATP concentration.

The approach we have used may be useful to study other complex proteins that undergo large conformational changes during their reaction cycle. *E. coli* RNA polymerase, for example, contains a 10 amino acid “trigger loop,” which covers the NTP binding site when the proper NTP is bound, apparently to hold it in place for addition to the growing RNA molecule. The tip of the trigger loop moves ∼20 Ǻ when the wrong, or no, NTP is present, apparently to allow only the correct NTP to occupy the active site for RNA synthesis ^43^. Mutational analysis of the corresponding part of *Saccharomyces cerevisiae* RNA polymerase II indicates that amino acids in the trigger loop interact with each other and with parts of the polymerase in other subunits, though close in 3D space and thus near the active site for RNA synthesis ^44^. We are not aware of mutational analyses of sites far from the active site of large enzymes akin to those reported here. We expect, however, that co-ordination of multiple activities of a large enzyme, such as RNA polymerase moving along the DNA template and adding correct NTPs, would require communication between distant parts of the enzyme, as discussed here for RecBCD. Kinesin and ATP synthase also undergo large conformational changes during their reaction cycles ^45-47^ and may be amenable to the type of analysis used here. RecBCD is large, has a complex reaction cycle, and is not essential for growth, making it readily amenable to mutational analysis. RecBCD is therefore an excellent model for studies in this area.

## Materials and Methods

### *E. coli* strains, growth media, and genetic methods

Media for cell growth, and methods for lambda and *E. coli* crosses and nuclease and Chi-cutting assays have been described ^40^. The *recC343* and *recB344* mutations ^28^ were cloned into pBR322 on the 19-kb *Bam*HI fragment containing *thyA-recC-ptrA-recB-recD-argA*, as described for *recBCD*^+ 30^. These plasmids, pDWS21 and pDWS33, respectively, were used for the genetic experiments described here and for DNA sequencing.

### Site-specific mutagenesis

Mutations were generated on plasmid pSA607, which contains the three genes of RecBCD controlled by their native promoters ^19^, using the QuikChange Site-Directed (Agilent Technologies) or the Q5 Site-Directed (New England Biolabs) mutagenesis kit. Oligonucleotides with the desired mutations were designed using web-based programs (Agilent Primer Design Program or NEBaseChanger) and purchased from Integrated DNA Technologies. Each mutant plasmid was confirmed by sequencing at least 300 bp on each side of the intended mutation, the only mutation detected. DNA primers for generating these mutations are available upon request. Plasmid pSA607 contains the *recC2773* mutation, which places six histidine residues on the RecC C-terminus to aid protein purification; this mutant has a wild-type phenotype ^19^. Allele numbers and codon changes are in Tables S1 – S3.

### Genetic assays for Chi hotspot activity and homologous recombination

Mutant plasmids were introduced by calcium-mediated transformation into strain V2831 (*ΔrecBCD2731<kan> argA21 recF143 hisG4 met rpsL31 galK2*); *ΔrecBCD2731* deletes the entire *recBCD* gene cluster ^41^. These transformants were tested for Chi hotspot activity, lambda recombination-proficiency, and *E. coli* recombination-proficiency using Hfr strain S727 (PO44 *thi-1 relA1*), as described ^18^.

### Enzymatic assays for general nuclease activity, DNA unwinding, and Chi cutting

RecBCD Chi-independent (“general”) nuclease activity was assayed using uniformly [^3^H]-labelled DNA from phage T7 ^33^. Duplicate assays, each with three amounts of extract in the linear range, are given in many cases in Table 2. Background values (*i.e.*, without ATP) determined in each set of assays were subtracted; values ranged from 21 to 58 units/mg of protein (37 ± 10 units/mg averaged over all assay sets). Protein concentrations were determined with a BioRad protein assay kit, using bovine serum albumin as standard. DNA unwinding and cutting at Chi were assayed using pBR322 with or without Chi (*χ*^+^*F225*), linearized with *Hin*dIII, and 5’-end labeled using [γ-^32^P] ATP and phage T4 polynucleotide kinase ^40^. Reaction products in Fig. 4 were separated by agarose gel electrophoresis and transferred to Whatman 3MM paper instead of DEAE paper; oligonucleotide limit-digestion products, seen in previously published results (e.g., ^40^) and Fig. S1 (top panel), were thus not detected in Fig. 4.

## Acknowledgments

For helpful discussions of protein structure, we are grateful to Rick Dahlquist, Brett Kaiser, Craig Kaplan, and Rick McLaughlin. We thank Randy Hyppa and Rasi Subramaniam for helpful comments on the manuscript and Dennis Schultz for plasmids pDWS21 and pDWS33. This research was supported by grants R35 GM118120 to G.R.S. and P30 CA015704 to the Fred Hutchinson Cancer Research Center from the National Institutes of Health of the United States of America.

## Author contributions

SKA, AFT, and GRS conceived the experiments, which SKA and AFT conducted. SKA, AFT, and GRS analyzed the data and wrote the manuscript.

## Additional information

Tables S1 – S3 contain recombination data for additional mutants in contact points CD, DB, and BC, respectively. Fig. S1 shows unwinding and Chi cutting activities of additional mutants. Fig. S2 shows contact point D4, close to B4 and C4. Fig. S3 shows the correlation of Chi hotspot activity and Hfr recombination proficiency for RecB tether mutants ^40^.

The authors declare no competing interests.

## Notes

### Competing Interest Statement

The authors have declared no competing interest.

### Summary of Updates

New data have been added, and the manuscript substantially rewritten.

