## Supplemental information for "Chi hotspot control of RecBCD helicase-nuclease by long-range intramolecular signaling"

SK Amundsen, AF Taylor, and GR Smith

**Supplementary Tables and Figures**

**Table S1. Mutants altered in RecC-RecD contact point (CD)**

| | | | | Genetic Assays | |
|---|---|---|---|---|---|
| Allele number[a] | Alternate allele designation[a] | C2[b] 541-544 QGEW | D2[b] 97-99 PTP | Chi hotspot activity[c] | *E. coli* Hfr cross (relative recombination frequency)[d] |
| WT | | . . . . | . . . | 5.1 ± 0.09 | 1.0 |
| *C2820* | C2△ | △△△△ | . . . | 1.5, 1.4 | 1.0, 0.64 |
| *C2821* | C2ala | AAAA | . . . | 3.2, 3.5 | 0.78, 0.67 |
| *D2824* | D2△ | . . . . | △△△ | 0.9, 1.0 | 0.43, 0.29 |
| *D2825* | D2ala | . . . . | AAA | 2.4, 2.9 | 0.38, 0.62 |
| *C2821 D2825* | C2ala D2ala | AAAA | AAA | 2.2, 2.3 | 0.66, 0.76 |
| *C2820 D2824* | C2△ D2△ | △△△△ | △△△ | 1.4, 1.1 | 0.81, 0.49 |

[a] The indicated mutation was on a derivative of plasmid pSA607 (*recBCD*$^+$) in strain V2831 (Δ*recBCD2731*). Alternate allele designations are those used in the main text.

[b] The contact point is composed of the indicated amino acids at the indicated positions. The alleles have mutations deleting the indicated amino acid (△); substitutions (when made) contain the indicated amino acid; (.) indicates that the amino acid is wild type.

[c] Chi hotspot activity in lambda vegetative crosses was determined as described in Materials and Methods [31]. Chi hotspot activity = $\sqrt{(t/c)_1/(t/c)_2}$ where t/c is the ratio of turbid ($cI^+$) to clear (*cI857*) recombinant plaques among $J^+$ $R^+$ recombinants in cross 1 (with $\chi^+$*D123*) and cross 2 (with $\chi^+$*76*).

[d] Frequency of His$^+$ [Str$^R$] recombinants per viable Hfr parent relative to that in the concurrent *recBCD*$^+$ cross. Wild-type frequency was 6.99% ± 0.46% (n = 25). The relative recombinant frequency for a null mutant (*recB21*) is 0.005.

**Table S2. Mutants altered in RecD-RecB contact point (DB)**

**Table S2A.**

| Allele number[a] | Alternate allele designation[a] | D3[b] 521-528 SVQPSRLP | Genetic assays | |
|---|---|---|---|---|
| | | | Chi hotspot activity[c] | *E. coli* Hfr cross (relative recombination frequency)[d] |
| WT | | ........ | 5.1 ± 0.09 | 1.0 |
| *D2826* | | ..E.DE.. | 3.5, 2.9 | 1.08, 0.94 |
| *D2827* | | ...A.... | 4.9, 4.4 | 1.06, 0.87 |
| *D2828* | | ......A. | 3.7, 3.2 | 0.76, 0.69 |
| *D2829* | | AA...... | 3.4, 2.8 | 0.58, 0.83 |
| *D2830* | | AA....A. | 1.5, 1.3 | 0.52, 0.39 |
| *D2831* | D3ala | ..AAAA.. | 1.9, 1.4 | 0.41, 0.49 |
| *D2832* | | AAAA.... | 2.8, 2.6 | 0.87, 0.76 |
| *D2833* | D3△ | ..△△△△.. | 1.0, 1.2 | 0.24, 0.24 |
| *D2834* | | ..△△.... | 4.0, 4.7 | 1.26, 0.87 |
| *D2835* | | ....△△.. | 5.1, 4.8 | 1.21, 0.69 |
| *D2836* | | △△...... | 0.96, 1.5 | 0.24, 0.46 |
| *D2837* | | ......△△ | 3.9, 3.7 | 0.79, 0.63 |
| *D2838* | | △△△△.... | 0.9, 1.1 | 0.39, 0.46 |
| *D2839* | | ..△△..△△ | 1.1, 1.2 | 0.23, 0.31 |
| *D2840* | | ....△△△△ | 0.9, 0.9 | 0.91, 0.79 |
| *D2841* | | △△..△△.. | 1.4, 1.2 | 0.31, 0.46 |
| *D2842* | | △....... | 3.1, 3.2 | 0.67, 0.77 |
| *D2843* | | .△...... | 1.6, 1.4 | 0.42, 0.61 |

Footnotes are as in Table S1.

**Table S2B.**

| Allele number[a] | Alternate allele designation[a] | B3[b] 634-646 DEHAWDVVVEEFD | Genetic assays | |
|---|---|---|---|---|
| | | | Chi hotspot activity[c] | *E. coli* Hfr cross (relative recombination frequency)[d] |
| WT | | ............. | 5.1 ± 0.09 | 1.0 |
| B2862 | | .A........... | 4.9, 4.8 | 0.87, 0.78 |
| B2863 | | .A...A....... | 4.1, 3.9 | 0.76, 0.72 |
| *B2864* | | .A...A...A... | 3.8, 3.9 | 0.84, 0.79 |
| *B2844* | | .A...A...A..A | 2.1, 2.2 | 0.31, 0.23 |
| *B2845* | B3ala | AA...A...AA.A | 1.8, 1.6 | 0.21, 0.34 |
| *B2846* | | .........Q..S | 3.6, 3.1 | 0.91, 0.78 |
| *B2847* | | .R...N...QK.S | 2.9, 3.0 | 0.83, 0.68 |
| *B2848* | | KR........... | 4.9, 4.8 | 0.92, 0.94 |
| *B2849* | | KR.......RR.. | 1.5, 2.1 | 0.48, 0.39 |
| *B2858* | | .........RR.. | 4.3, 3.9 | 0.39, 0.54 |
| *B2850* | | KR.......RR.K | 1.7, 1.5 | 0.42, 0.33 |
| *B2851* | B3lys,arg | KR...K...RR.. | 1.2, 1.4 | 0.39, 0.42 |
| *B2852* | B3△ | △△△△△△△△△△△△△ | 0.9, 1.0 | 0.006, 0.009 |
| *B2853* | | .....△△△△△... | 4.1, 4.7 | 0.87, 0.93 |
| *B2854* | | △△△△△........ | 4.9, 3.5 | 0.93, 0.68 |
| *B2855* | | ..........△△△ | 3.8, 3.6 | 0.29, 0.38 |
| *B2856* | B3△(10) | △△△△△△△△△△... | 4.6, 4.2 | 0.88, 0.78 |
| *B2857* | | .....△△△△△△△△ | 3.1, 2.7 | 0.08, 0.07 |
| *B2859* | | .........AAAA | 4.8, 4.9 | 0.68, 0.84 |

**Table S2C.**

| Allele number[a] | Alternate allele designation[a] | B3[b] 634-646 DEHAWDVVVEEFD | D3[b] 521-528 SVQPSRLP | Genetic Assays | |
|---|---|---|---|---|---|
| | | | | Chi hotspot activity[c] | *E. coli* Hfr cross (relative recombination frequency)[d] |
| WT | | ............. | ........ | 5.1 ± 0.09 | 1.0 |
| *B2845 D2830* | B3ala D3ala | AA...A...AA.A | AA....A. | 1.5, 1.4 | 0.48, 0.39 |
| *B2845 D2833* | B3ala D3△ | AA...A...AA.A | ..△△△△.. | 1.1 ± 0.06 | 0.53, 0.34 |
| *B2844 D2833* | | .A...A...A..A | ..△△△△.. | 1.8, 1.6 | 0.42, 0.38 |
| *B2847 D2826* | | .R...N...QK.S | ..E.DE.. | 1.8 ± 0.2 | 0.66 ± 0.1 |

Footnotes are as in Table S1.

**Table S3. Mutants altered in or near RecB-RecC contact point (BC)**

| Allele number[a] | Alternate allele designation[a] | B4[b] 913-922 GHGIAQDLMP | C4[b] 599-608 FLPDAETEAA | D4[b] 469-475 HRHPHSR | Genetic assays: Chi hotspot activity[c] | Genetic assays: *E. coli* Hfr cross (relative recombination frequency[d] |
|---|---|---|---|---|---|---|
| WT | | .......... | .......... | ....... | 5.1 ± 0.09 | 1.0 |
| *B2865* | B4ala | AAAAAAAAAA | .......... | ....... | 3.6, 3.7 | 0.58, 0.41 |
| *B2860* | B4△ | △△△△△△△△△△ | .......... | ....... | 4.2, 4.3 | 0.81, 0.52 |
| *C2861* | | .......... | ...A.A.A.. | ....... | 4.9, 3.8 | 0.87, 0.68 |
| *C2823* | C4△(7) | .......... | ...△△△△△△△ | ....... | 2.3 ± 0.12 | 0.32 ± 0.03 |
| *C2822* | C4△ | .......... | △△△△△△△△△△ | ....... | 1.1 ± 0.12 | 0.24 ± 0.04 |
| *D2867* | | .......... | .......... | AAAAAAA | 4.4 ± 0.15 | 0.74 ± 0.06 |
| *D2868* | | .......... | .......... | △△△△△△△ | 3.9 ± 0.17 | 0.83 ± 0.07 |
| *B2860* | B4Δ | △△△△△△△△△△ | .......... | ....... | 4.2, 4.3 | 0.81, 0.52 |
| *B2860 D2868* | | △△△△△△△△△△ | .......... | △△△△△△△ | 5.2, 4.3 | 1.31, 0.59 |
| *B2865 D2868* | | AAAAAAAAAA | .......... | △△△△△△△ | 4.0, 4.6 | 0.22, 0.29 |
| *C2822* | C4Δ | .......... | △△△△△△△△△△ | ....... | 1.1 ± 0.12 | 0.24 ± 0.04 |
| *C2823* | C4Δ(7) | .......... | ...△△△△△△△ | ....... | 2.3 ± 0.12 | 0.32 ±0.03 |
| *B2865 C2861* | | AAAAAAAAAA | ...A.A.A.. | ....... | 2.5 ± 0.26 | 0.31 ± 0.09 |
| *C2861 D2868* | | .......... | ...A.A.A.. | △△△△△△△ | 3.9 ± 0.14 | 0.53 ± 0.08 |
| *B2860 C2822* | B4Δ C4Δ | △△△△△△△△△△ | △△△△△△△△△△ | ....... | 1.1, 1.3 | 0.09, 0.22 |
| *B2865 C2861 D2868* | | AAAAAAAAAA | ...A.A.A.. | △△△△△△△ | 3.4 ± 0.27 | 0.44 ± 0.06 |

Footnotes are as in Table S1.

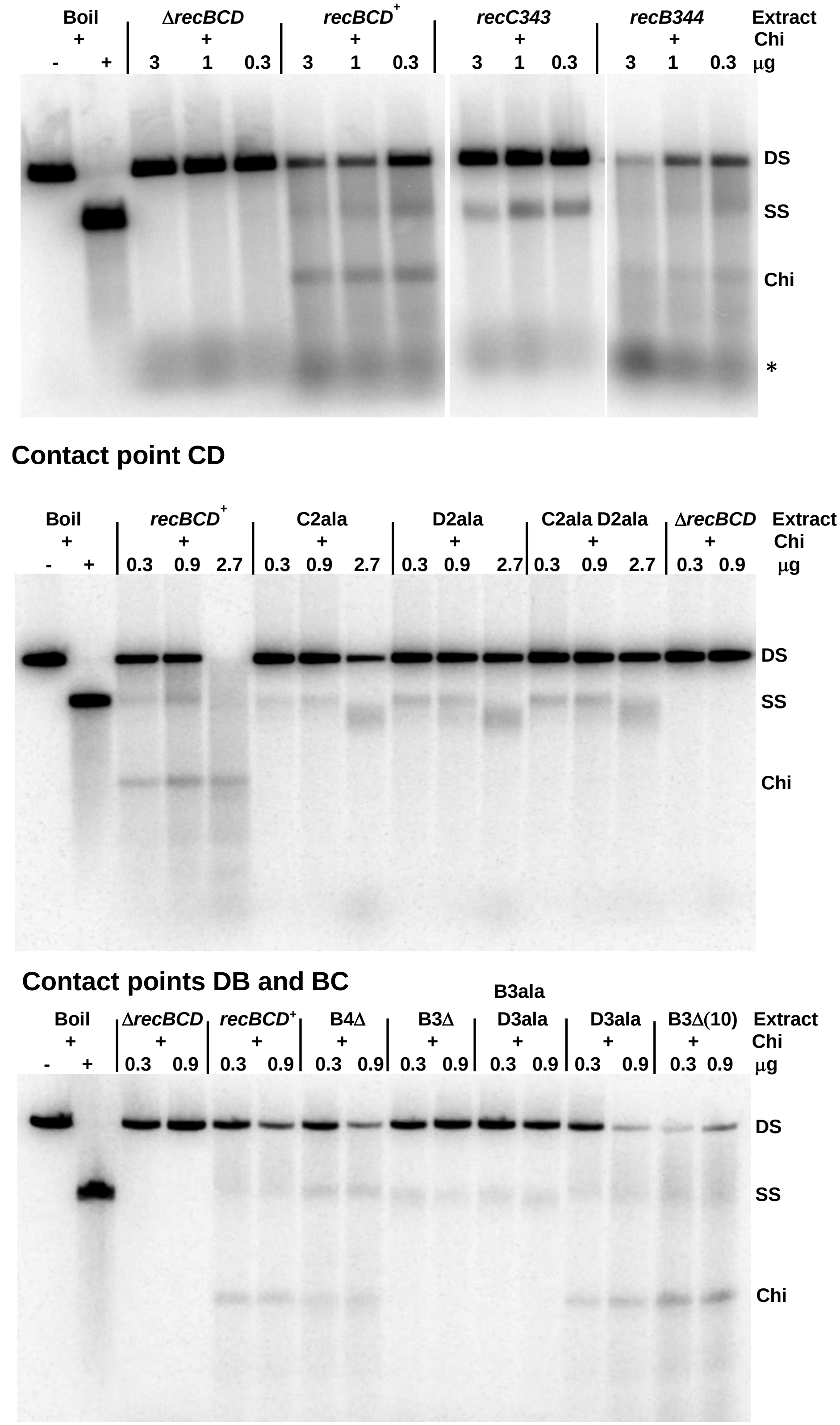


**Figure S1. RecBCD contact-point mutants, *recC343,* and *recB344* retain DNA unwinding activity but have reduced or undetectable cutting of DNA at Chi hotspots**

Extracts of the indicated mutants were assayed for unwinding and cutting of linear pBR322 DNA (4.3 kb long) with a Chi site ($\chi^{+}$*F225*) 1470 bp from the 5' [$^{32}$P]-labelled DNA end. Note that the Chi-cut species and ss DNA are reaction intermediates and their observed amount is not necessarily a linear function of enzyme amount. ds substrate (DS), unwound ss DNA (SS), Chi-cut DNA (Chi) and limit digestion products (oligonucleotides; *) are indicated.

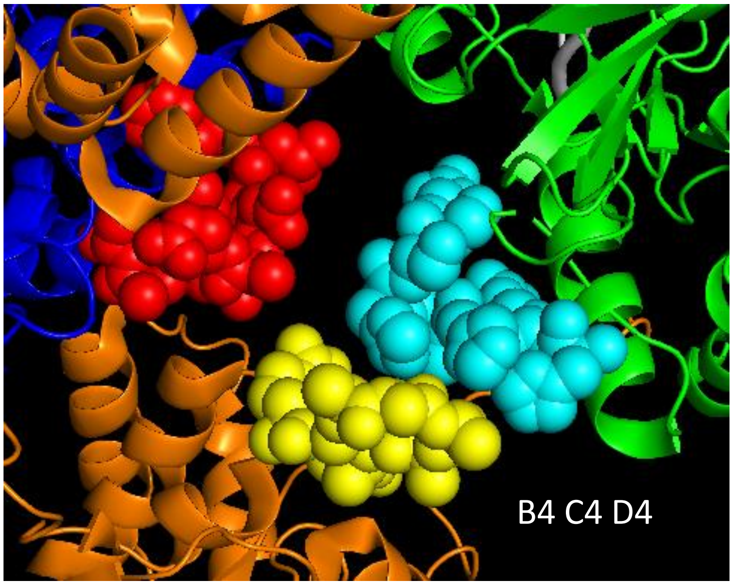


**Figure S2. Contact point D4, close to contact points B4 and C4.** Contact points are shown as spheres. B4 (amino acids GHGIAQDLMP at positions 913 – 922, modeled as alanines, of RecB) is yellow, C4 (amino acids FLPDAETEAA at positions 599 – 608 of RecC) is red, and D4 (amino acids HRHPHSR at positions 469 – 475 of RecD) is cyan. Shown is part of the cryoEM structure PDB 5LD2.

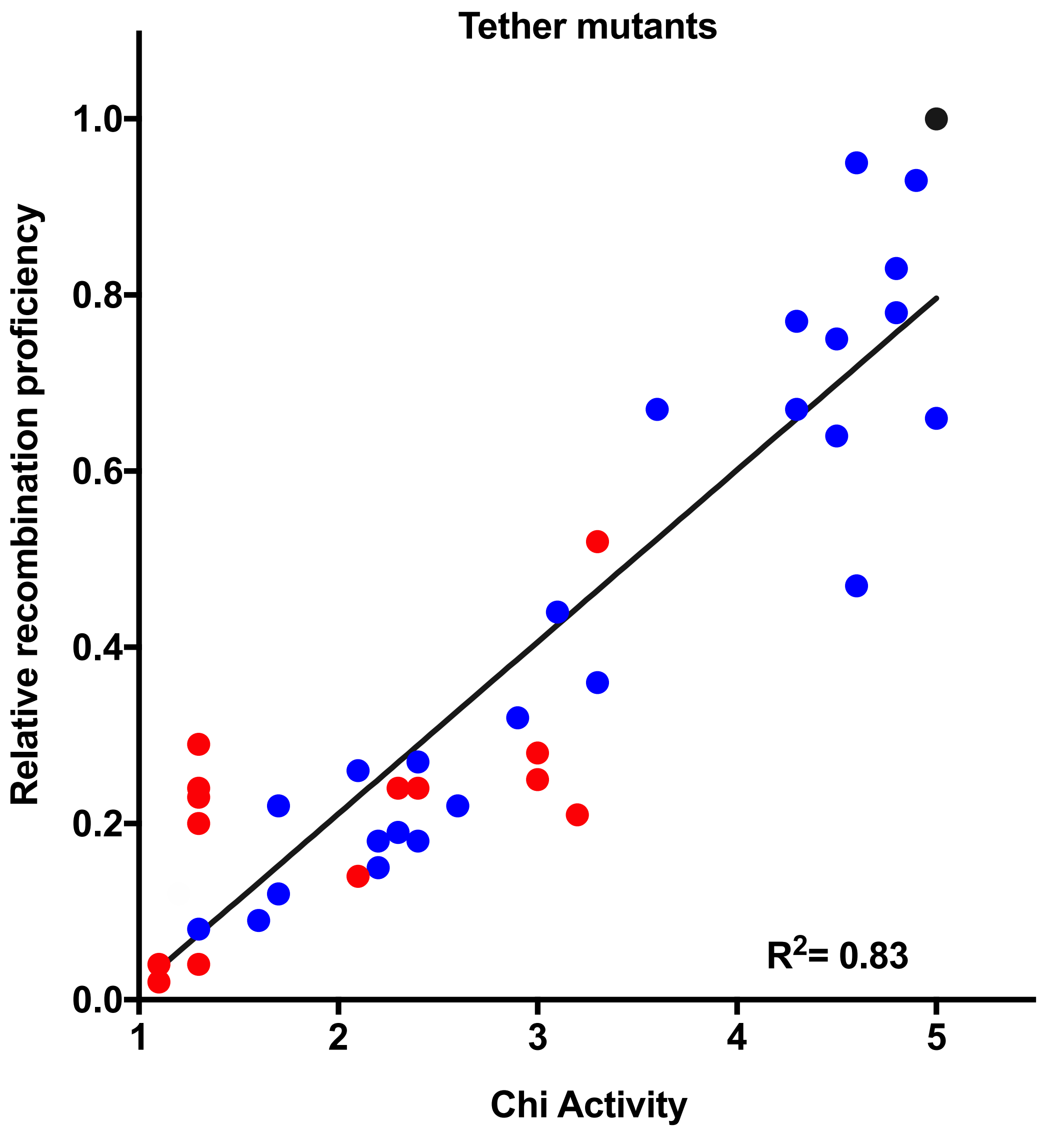


**Fig. S3.** ***E. coli*** **Hfr recombination proficiency is positively correlated with Chi hotspot activity in RecC tunnel mutants.** Compare with Figure 3 of the main text. Red data points are for deletion mutants and blue for substitution mutants; black point is wild type. $R^2 = 0.83$ ($p < 0.0001$). Modified from [40].

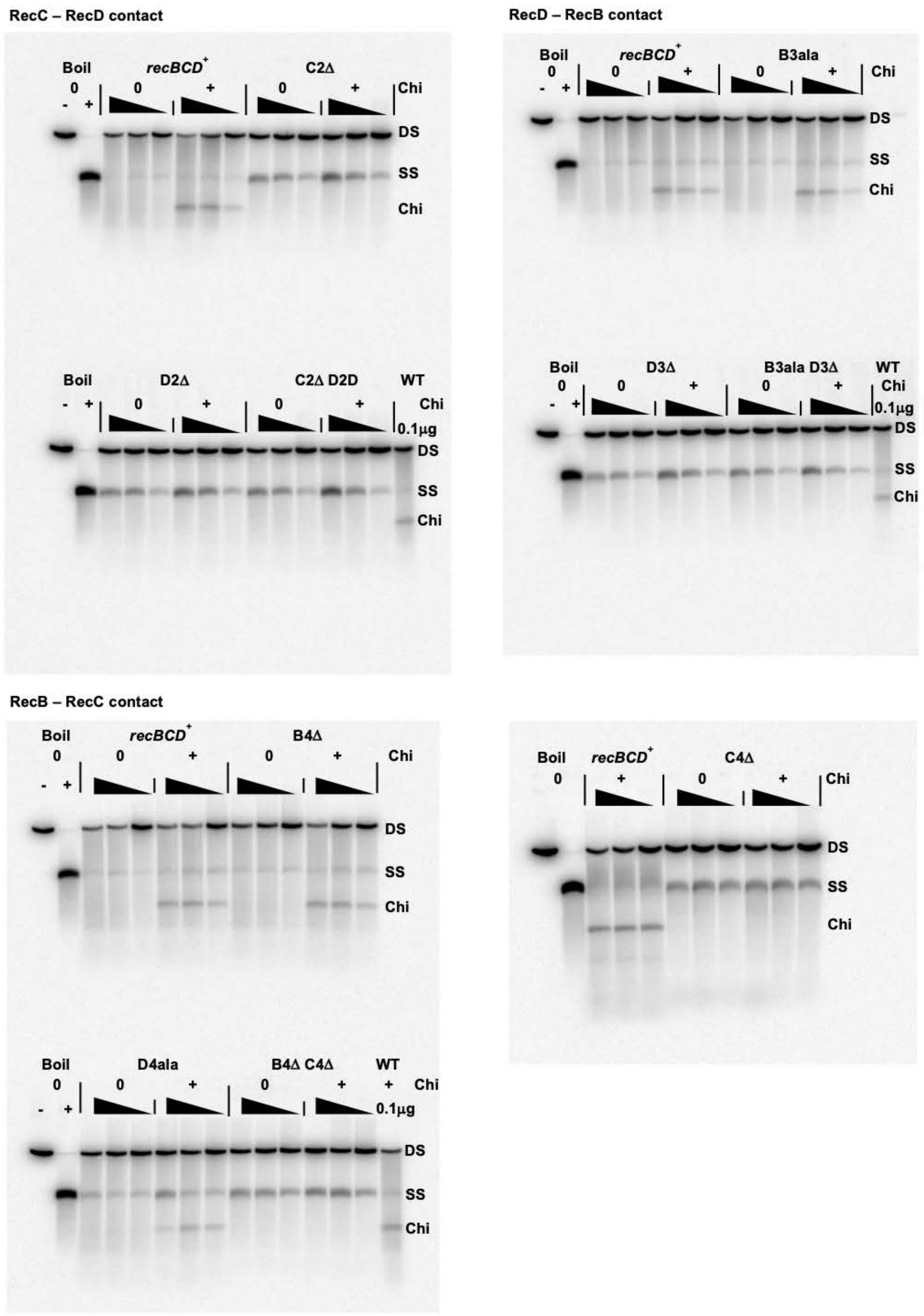


**Supplementary Figure S4. RecBCD contact-point mutants retain DNA unwinding but have reduced or undetectable cutting of DNA at Chi hotspots**. Shown are full-length blots of the gels corresponding to the data in Figure 4.

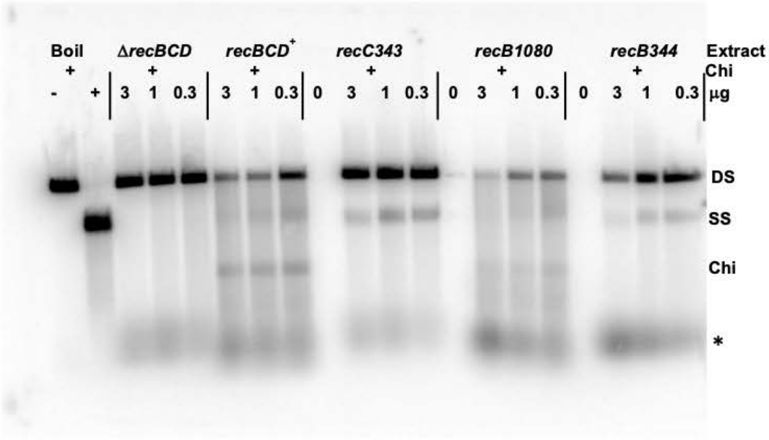


**Contact point CD**

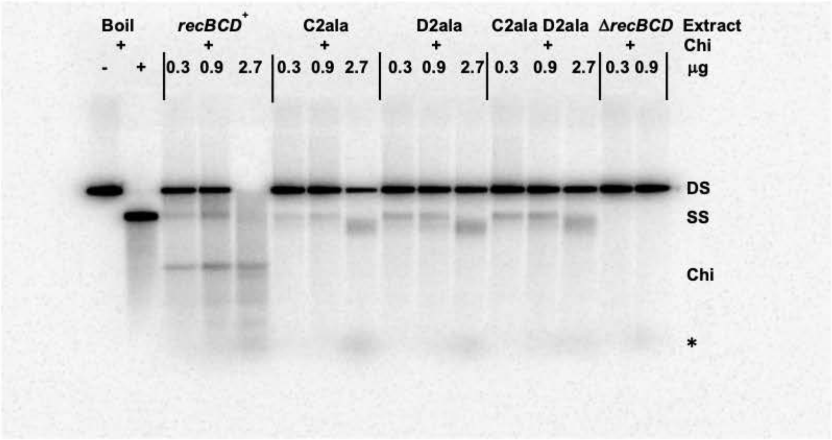


**Contact points DB and BC**

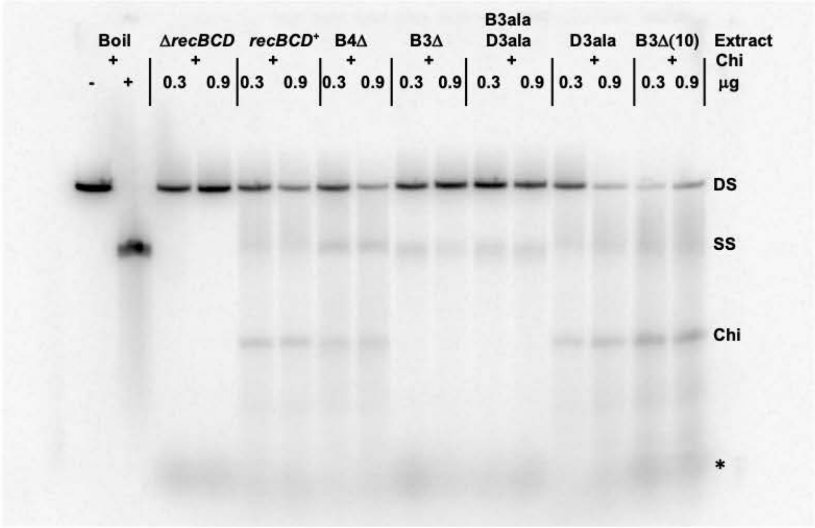


**Supplementary Figure S5. RecBCD contact-point mutants, *recC343,* and *recB344* retain DNA unwinding activity but have reduced or undetectable cutting of DNA at Chi hotspots**
Shown are full-length blots of the gels corresponding to the data in Figure S1.
